# An efficient method to clone TAL effector genes from *Xanthomonas oryzae* using Gibson assembly

**DOI:** 10.1101/723064

**Authors:** Chenhao Li, Chonghui Ji, José C. Huguet-Tapia, Frank F. White, Hansong Dong, Bing Yang

## Abstract

TALes (Transcription Activator-Like effectors) represent the largest family of type III effectors among pathogenic bacteria and play a critical role in the process of infection. Strains of *Xanthomonas oryzae* pv. oryzae (Xoo) and some strains of other *Xanthomonas* pathogens contain large numbers of TALe genes. Previous techniques to clone individual or a complement of TALe genes through conventional strategies are inefficient and time-consuming due to multiple genes (up to 29 copies) in a given genome and technically challenging due to the repetitive sequences (up to 33 nearly identical 102-nucleotide repeats) of individual TALe genes. Thus, only a limited number of TALe genes have been molecularly cloned and characterized, and the functions of most TALe genes remain unknown. Here, we present an easy and efficient cloning technique to clone TALe genes selectively through *in vitro* homologous recombination and single strand annealing and demonstrate the feasibility of this approach with four different Xoo strains. Based on the Gibson assembly strategy, two complementary vectors with scaffolds that can preferentially capture all TALe genes from a pool of genomic fragments were designed. Both vector systems enabled cloning of a full complement of TALe genes from each of four Xoo strains and functional analysis of individual TALes in rice in approximately one month compared to three months by previously used methods. The results demonstrate a robust tool to advance TALe biology and a potential for broad usage of this approach to clone multiple copies of highly competitive DNA elements in any genome of interest.

## INTRODUCTION

Phytopathogenic bacteria of the genus *Xanthomonas* cause a variety of plant diseases including those inflicting severe losses on the economically important crop plants worldwide (Leyns et al., 1984). In rice, *Xanthomonas oryzae* pv. *oryzae* (Xoo) invades xylem to cause bacterial blight of vascular disease, while the closely related pathogen *X. oryzae* pv. *oryzicola* (Xoc) is a mesophyll colonizer and causes a disease known as bacterial leaf streak (Niño-Liu et al., 2006). Individual strains of both Xoo and Xoc contain multiple (e.g., up to 29 in a typical Xoc strain) genes of the Transcription Activator-Like effector (TALe) family, a few of which are known virulence or/and avirulence effectors depending on the genetic context of host plants (White, 2016). TALes depend on a type III secretion system for their translocation to host cells (Zhang et al., 2015). Typically, once internalized into nuclei of rice cells, TALes act upon the effector binding elements (EBEs) in the promoters of host genes (Boch et al., 2010). The general hypothesis is that TALes have evolved to target specific host genes, whose subsequent ectopic expression facilitates infection, and, indeed, a number of TALe-targeted genes have been shown to enhance host susceptibility (Chakrabarty et al., 1997; Yang et al., 2006; Sugio et al., 2007; Yu et al., 2011; Bart et al., 2012; Cohn et al., 2014; Hu et al., 2014; Cox et al., 2017; Schwartz et al., 2017). However, relatively few TALe targets have been shown to be disease susceptibility (S) genes given the large number of TALe genes discovered. Plants have also evolved to turn TALe function against bacterial invasion, and a few TALes trigger host resistance response due to TALe-mediated expression of the so-called executor resistance (R) genes, which, in rice, includes*Xa27, Xa10* and *Xa23* (Gu et al., 2005; Tian et al., 2014; Wang et al., 2015). TALes are also recognized by members of the common NBS LRR family of R genes (e.g., *Xa1*) (Ji et al., 2016). A group of TALe variants or iTALes (interfering TALes or truncated TALes) from Xoo and Xoc can suppress NBS LRR-mediated TALe recognition and resistance (Ji et al., 2016; Read et al., 2016). However, additional machenisms by which a large number of TALes are involved in host/pathogen interaction remain largely unexplored, which requires a simple and efficient way to clone and characterize those TALe genes.

TALe genes encode three major functional domains. The N-terminal domain contains signal for bacterial type III secretion, followed by the central tandem repetitive region, which specifies the target nucleotide sequence of host genes, and the C-terminal domain containing nuclear localization signals and transcription activation domain, the latter characteristic of features of eukaryotic transcription factors (Boch et al., 2010). The specificity of interaction between the TALes and their host target DNA is determined by the combination of number of central repeats and composition of two amino acids at the 12^th^ and 13^th^ positions of repeats of TALes known as the repeat variable diamino acid or RVD unit (Boch et al., 2009; Moscou and Bogdanove, 2009). Coding sequences of TALes contain the *Sph*I or *Bam*HI restriction sites flanking the central or almost whole region across different *Xanthomonas* subspecies or pathovars are highly conserved. Often, the native TALe genes were not cloned, rather the repeat domains were cloned using the conserved *Sph*I or *Bam*HI sites of both ends of TALe genes. The cloned *Sph*I or *Bam*HI fragment fused with a common scaffold of sequences for N-terminal and C-terminal domains of TALe bestows specificity of the new TALe gene (Hopkins et al., 1992; Yang and White, 2004). The characterization of TALe genes from various *Xanthomonas* strains has primarily involved cloning by library construction followed by hybridization or PCR detection based on a conserved TALe sequence (Leach et al., 1992; De Feyter et al., 1993; Yang and White, 2004; Yu et al., 2015; Tran et al., 2018b). Cloning and screening for the right TALe genes are technically challenging due to existence of many copies of TALe genes in a genome of *Xanthomonas* and excessive near-identical tandem repeats in individual TALe genes. Sequencing through the whole central repetitive region of TALe gene is difficult. The whole cloning process is also time-consuming and inefficient, taking more than 3 months to clone and test a TALe gene from a given *Xanthomonas* strain.

To improve the efficiency of cloning TALe genes from complex genomes, in present study, we adapted the Gibson assembly strategy using the conserved nature of TALe genes. The Gibson assembly method is a robust molecular cloning technology alternative to restriction/ligation subcloning (Gibson et al., 2009). The method depends on an exonuclease to excise the 5’ nucleotides of double stranded DNA (dsDNA) to produce a 3’ single strand overhang, which allows complementarity or annealing to the single strand overhang of an adjacent fragment also caused by the exonuclease. A DNA polymerase extends the nucleotides at the 3’ overhangs, and a DNA ligase is used to seal the nicks. The three reactions can be executed in a single tube using a programmed protocol in a thermocycler, and the reaction can be transferred directly into host cells for replication of the plasmids (Gibson et al., 2009). The strategy was applied to selectively clone the full complements of TALe genes from four strains of Xoo, representing the Asian and African lineages of the pathogen. The resulting TALe genes were then assessed for their abilities to restore virulence to a TALe-defective mutant of Xoo.

## RESULTS

### Similarity of TAL effector genes in *Xanthomonas*

Gibson assembly requires the overlapping sequences of approximately 20 bp between two adjacent DNA fragments, and the process is summarized in Figure 1. The TALe gene sequences were retrieved from genome data of thirty-three Xoo, ten Xoc, two *X. citri* pv. *vignicola* and one *X. citri* pv. *malvacearum* strains in the NCBI (National Center for Biotechnology Information) database (Supplementary Table S1). TALe gene content of twenty-two Asian Xoo strains and eleven African Xoo strains ranges from 10 to 20 and 9, respectively, in each strain. Xoc strains contain a range of 20-29 TALe genes. Most TALe genes contain two *Bam*HI restriction sites (GGATCC), one at the ATG start codon and another approximately 150 bp upstream of the stop codon, and two *Sph*I sites (GCATGC), one 33 bp upstream of the first repeat and the second one located about 450 bp downstream of the last repeat. Sequences around these restriction sites are also very conserved, almost 99% identical. However, some rare TALe genes contain single nucleotide polymorphism (SNP) within one of the *Sph*I or *Bam*HI recognition sequences. Only 40 out of 733 TALe genes (5.4%) contained a 1-bp variation in one of the two *Bam*HI restriction sites (GGATCC to GGATCT). Fifty-seven of 733 TALe genes (7.8%) contain a SNP within one of the *Sph*I restriction sequences (Suppl. Table 1). The loss of either *Bam*HI or *Sph*I sites prevents cloning of *Bam*HI or *Sph*I fragment of corresponding TALe gene in a library made from either *Bam*HI or *Sph*I DNA fragments. The results indicate that the majority (>92%) of TALe genes can be cloned through capture of either *Bam*HI or *Sph*I fragments of full-length genes (Supplementary Table S1), and rare TALe gene variants can be retrieved using other restriction enzyme combinations.

**Fig. 1.**
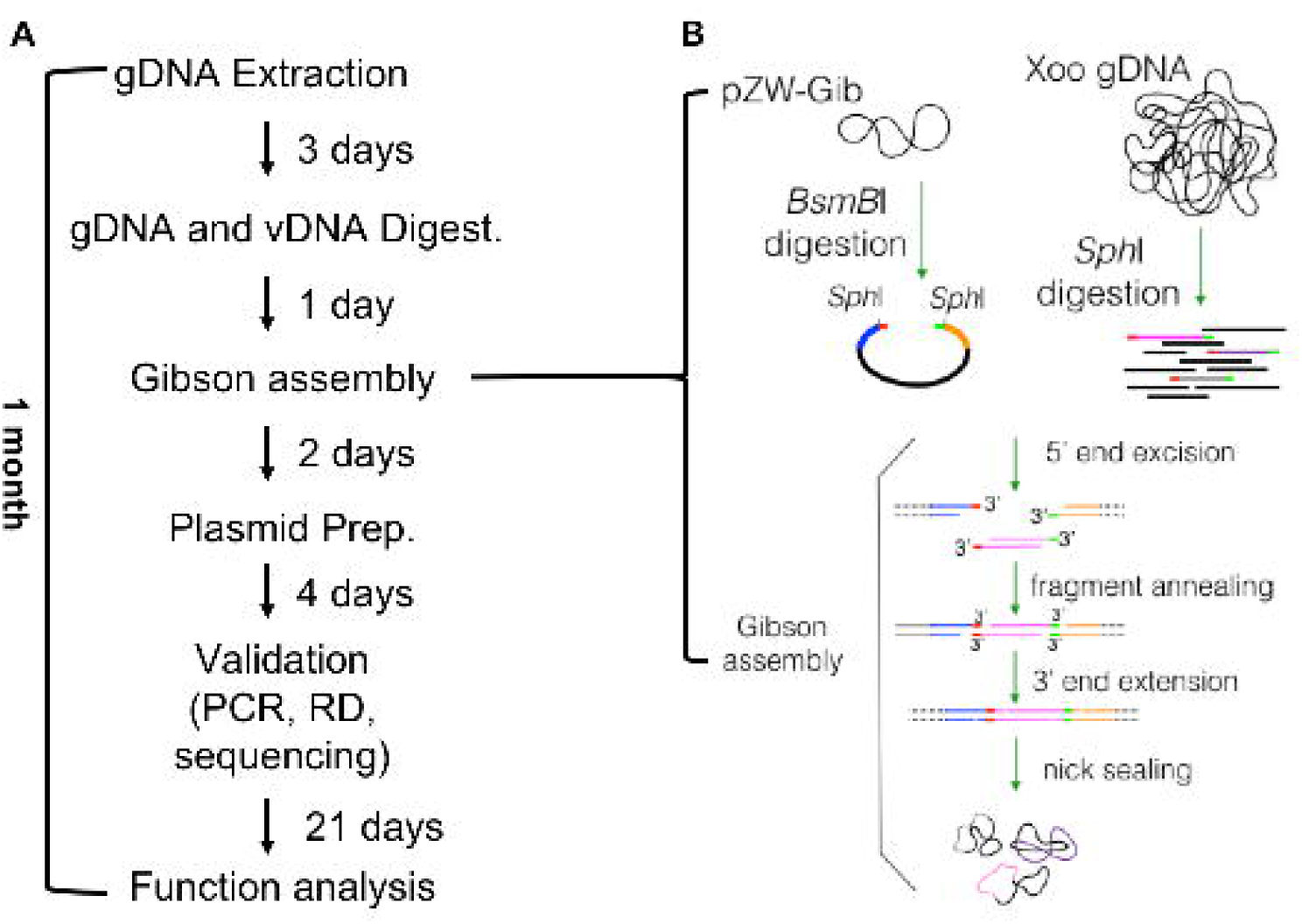
The flow chart and schematics of selectively isolating TALe genes using Gibson assembly method. **A.** Steps and timing of cloning and analysis of TALe genes from a given Xoo genome. **B.** The selective cloning of *Sph*I fragments of TALe genes. *Bsm*BI digestion yields cloning vector containing a homologous end (hook, colored end) at each side beyond *Sph*I site; *Sph*I digestion of genomic DNA of Xoo releases TALe fragments with both ends containing short sequences complementary to the hooks of cloning vector pZW-Gib. Gibson assembly results in plasmids containing individual TALe genes.

### Strategy to selectively clone TALe genes

We next developed a system to selectively isolate the genomic fragments, e.g., *Sph*I or *Bam*HI digested fragments, of TALe genes from a genome. The technique involves *in vitro* linking a TALe fragment and a vector fragment with a short (∼20 bp) 5’ and 3’ end sequences matching to both ends of TALe gene fragments by annealing of the 3’ complementary overhangs produced by T5 exonuclease. The single stranded overhangs of the vector fragment will function as sequence-specific hooks to catch the corresponding overhangs of TALe fragments selectively from a pool of the genomic fragments. To make a cloning vector for selectively cloning of the *Sph*I fragments of TALe genes, the previously cloned TALe gene *pthXo1*, which encodes the major virulence effector for the Asian Xoo strain PXO99^A^ (Yang and White, 2004), was used as the backbone of TALe gene vector without the central *Sph*I repetitive region. In addition, a counter selective gene was introduced to avoid undigested cloning vector or vector insert. First, a DNA fragment (gBlock) was synthesized that contained the *ccdB* gene, flanked by a sequence (5’-<u>GCATGC</u>ATGGCGCAATGCACTGACGGGTGCA<u>GAGACG</u>-3’), 23 nucleotides (nt) downstream of the first *Sph*I (underlined), and a sequence (5’-<u>CGTCTC</u>AACGCCGGATCAGGCGTCTTT<u>GCATGC</u>-3’), 20 nucleotides upstream of the second *Sph*I of *pthXo1*. The two *ccdB* flanking sequences between *Sph*I and *Bsm*BI (double underlined) sites are highly conservative among the natural TALe genes (Supplementary Fig. S1A). The *ccdB* gene encodes a toxic protein (CcdB) to cause cell death in certain *E. coli* strains (Bernard and Couturier, 1992). Inclusion of *ccdB* insures loss of any clones that retains the region of *ccdB* when transferred to a strain lacking the antidote *ccdA* gene, in this case *E. coil* XL1-Blue. Digestion of vector with *Bsm*BI releases two ends (23 and 20 nt) that are homologous to the two ends of the *Sph*I fragment of TALe gene. The gBlock was cloned into pZW-pthXo1 at *Sph*I sites by replacing the central *Sph*I region of *pthXo1*, resulting in pZW-Gib. The backbone of TALe in pZW-Gib contains the conserved ∼800 bp region upstream and ∼400 bp region downstream of the *Sph*I recognition sites of TALe genes (Fig. 2A). Insertion of the *Sph*I central repeat region of any TALe gene through Gibson assembly should result in functional TALe genes.

**Fig. 2.**
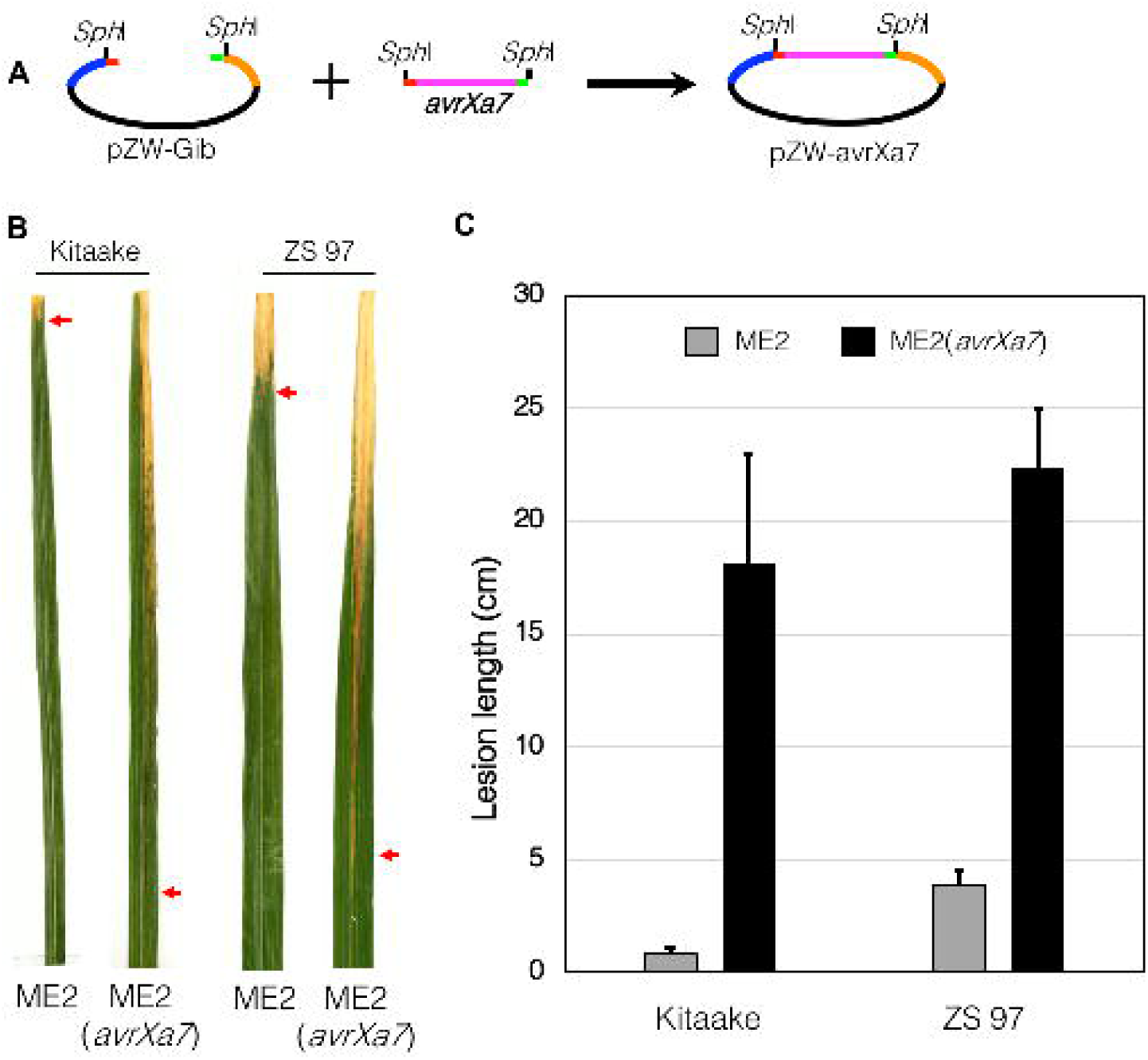
Validation of pZW-Gib for cloning of a functional TALe gene. A. The *Sph*I fragment of *avrXa7* was cloned into pZW-Gib, resulting in pZW-avrXa7. **B**. Rice leaves showing disease symptoms caused by Xoo strains. **C**. Lesion lengths in rice leaves caused by Xoo strains ME2 and its transformant ME2(*avrXa7*). Ten fully expanded young leaves were inoculated for lesion length measurements 14 days post inoculation. The experiment was repeated twice independently with similar results.

To test the system, pZW-Gib was digested with *Bsm*BI to remove the *ccdB* gene and used to clone the *Sph*I fragment of the alternate major virulence TALe gene *avrXa7*, which was originally cloned from the Xoo strain PXO86 (Fig. 2A). The pZW-Gib-avrXa7 was introduced via subcloning into pHM1, a Xoo suitable plasmid, and transferred to ME2, a mutant derived from PXO99^A^ with *pthXo1* inactivated (Yang and White, 2004). Finally, the function of *avrXa7* for virulence, as assembled in pZW-Gib, was tested in the rice cultivars Kitaake and Zhenshan 97. The results showed that the Gibson-cloned *avrXa7* restored the virulence in ME2(*avrXa7*) in term of lesion lengths and disease symptoms compared to ME2 (Fig. 2B; 2C).

### Isolation and functional test of TALe genes from PXO61

The PXO61 genome of Xoo (NCBI accession, CP033187.1) has over 3,000 *Sph*I sites and eighteen TALe genes (Supplementary Fig. S3A). All TALe genes contain two *Sph*I sites except one, *Tal5b*, which contains only one *Sph*I restriction site with the second one missing due to a deletion in the 3’ region. *Tal5b* belongs to a group of iTALe genes that contribute virulence to Xoo by suppressing *Xa1* mediated disease resistance (Ji et al., 2016). After ligation, the *Sph*I fragments of genomic DNA derived from PXO61 into the *Bsm*BI-digested pZW-Gib and, using the Gibson assembly protocol, bacterial colonies were picked randomly to screen for the presence of *Sph*I fragments of TALe genes (Fig. 3A, 3B). Twenty-two out of twenty-six clones were positive for TALe genes based on PCR with specific primers, restriction enzyme digestion and Sanger sequencing (Fig. 3B). To clone the whole complement of TALe genes (n=17 except *Tal5b*) of PXO61, a total of 45 clones were selected for sequencing based on the sizes of *Sph*I fragments of clones and the predicted number of TALe genes in the PXO61 genome. Seventeen of the cloned TALe genes matched the corresponding annotated genes in PXO61 genome (Fig. 4). The individual TALe genes in pZW-Gib were inserted into the broad host range vector pHM1, and the constructs transferred to ME2 for functional analysis. One clone contained the previously identified major TALe gene *pthXo3* (also known as *Tal6C*), which conferred disease susceptibility in rice Kitaake and Zhenshan 97 (Fig. 5, Supplementary Fig. S4C), and another clone contained *Tal7* (now *pthXo2B*), encoding a virulence TALe targeting the S gene *SWEET13* and confers disease susceptibility only to Kitaake and not Zhenshan 97 (Fig. 5, Supplementary Fig. S4C). The remaining 15 clones conferred no observable phenotype (Fig. 5; Supplementary Fig. S4C).

**Fig. 3.**
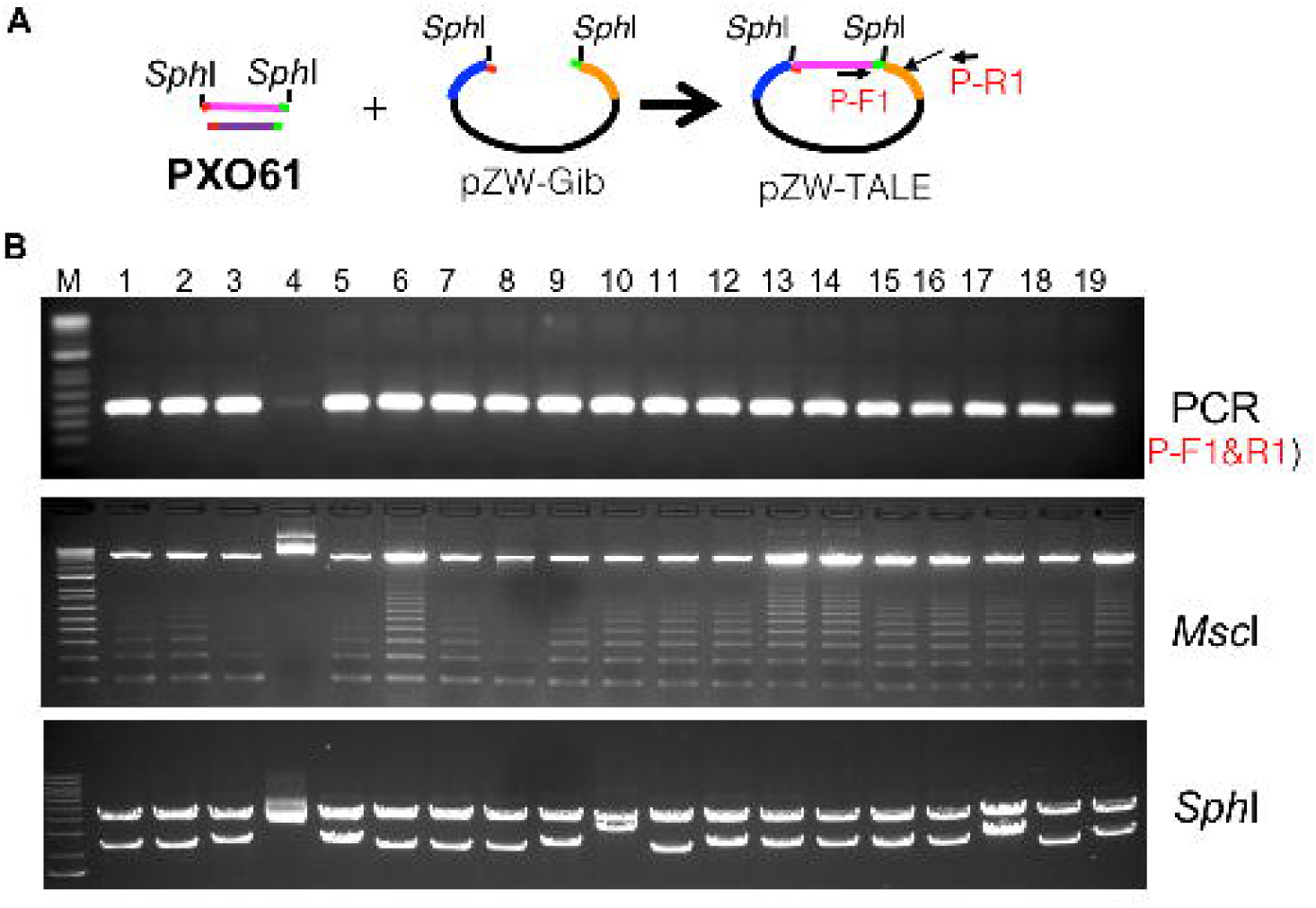
Validation of TALe clones through PCR, restriction enzyme digestions. **A.** Schematic of selective isolation of *Sph*I fragments TALe genes from genomic DNA of PXO61. **B.** Validation of TALe clones through PCR with primers P-F1 and P-R1 of individual clones as indicated above lanes of upper gel image, digestion by *Msc*I which cuts each of central repeats (the DNA band patterns resulted from partial digestion), and digestion by *Sph*I.

**Fig. 4.**
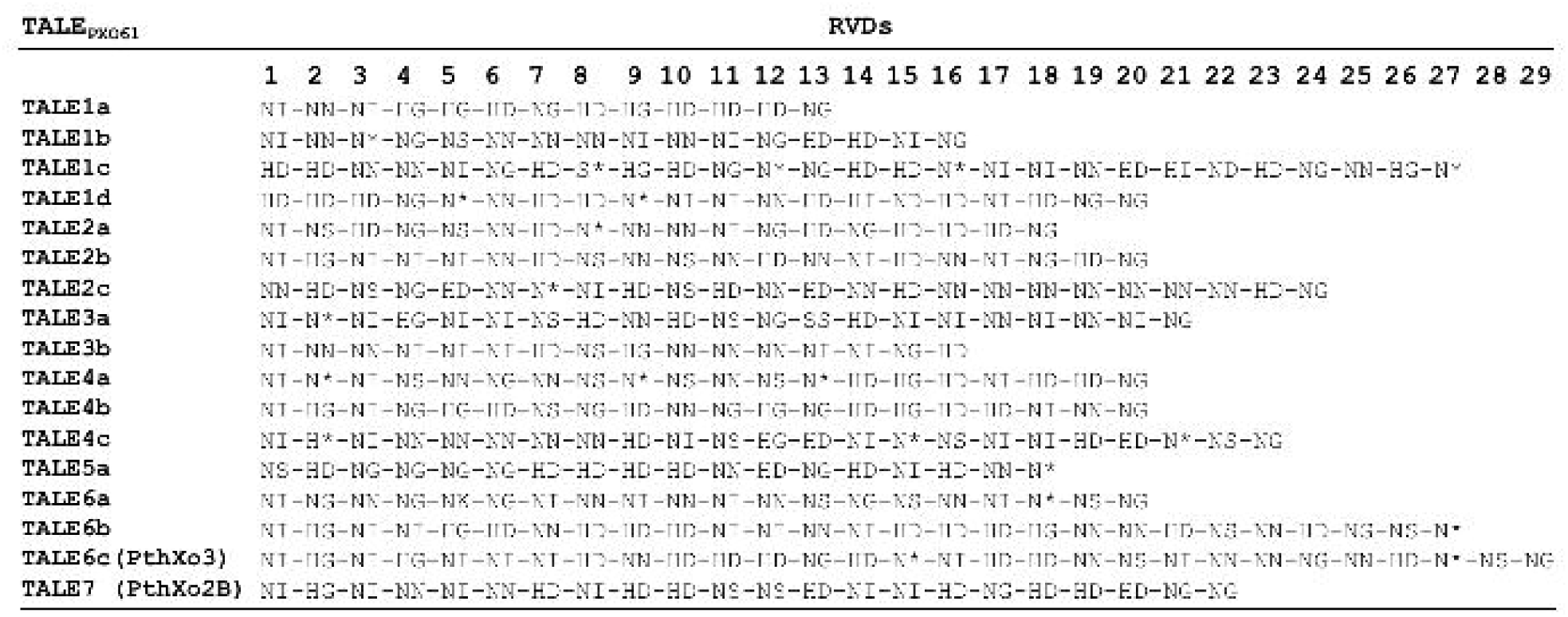
Seventeen of eighteen annotated TALe genes from PXO61 were cloned using pZW-Gib vector and Gibson assembly method. The RVDs of individual TALes are shown under numbers (1 to 29) indicating the order of 33-34 amino acid repeats. Asterisk (*) indicates amino acid at the 13^th^ position missing.

**Fig. 5.**
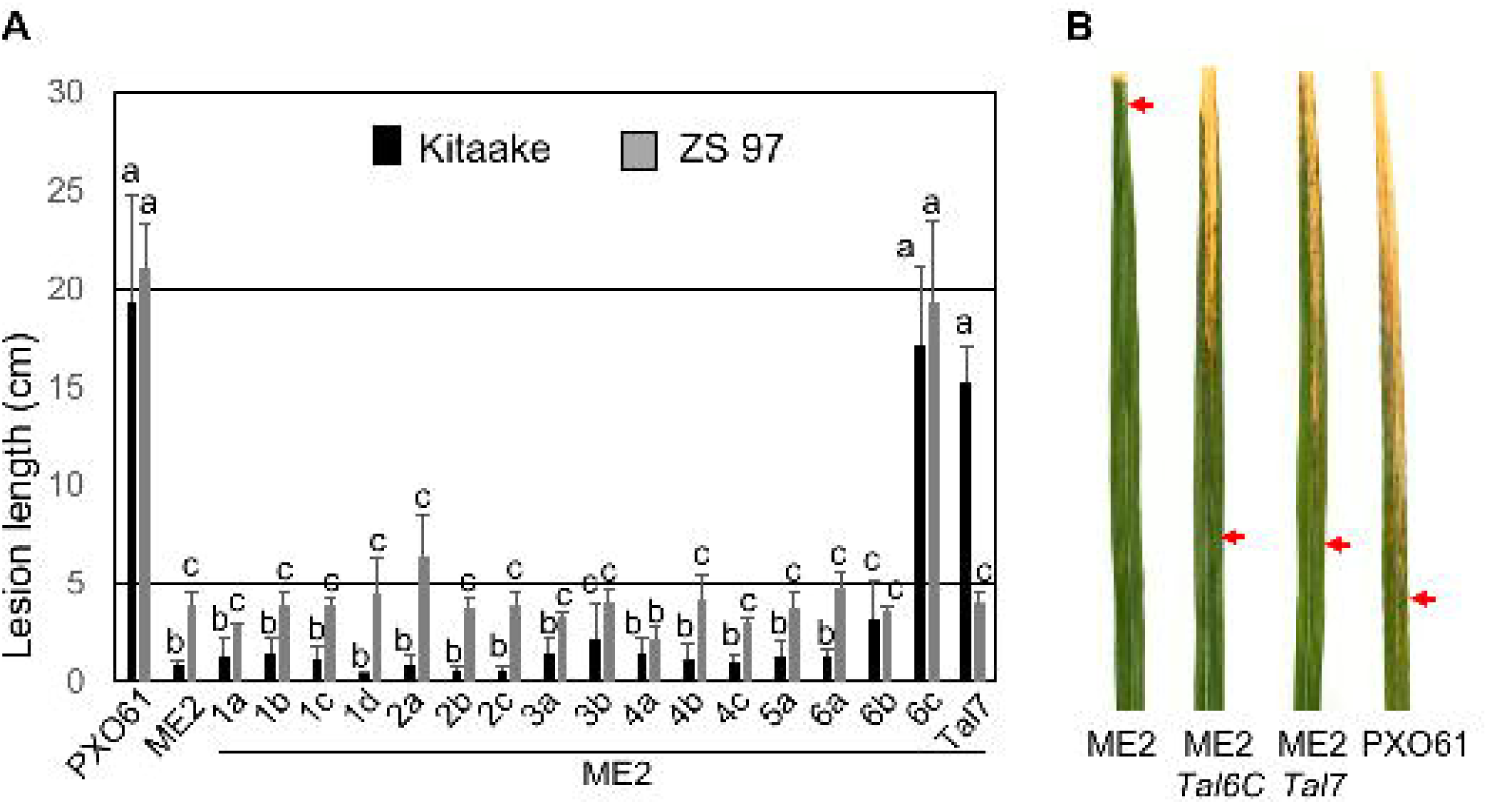
Virulence contribution of seventeen TALes cloned from PXO61. **A.** Lesion lengths caused in rice Kitaake and Zhenshan 97 (ZS 97) by different Xoo strains indicated below the paired columns. Different lower letters indicate statistically significant difference (means ± s.e.m, n=10, *P*<0.05). **B.** Blight symptom in Kitaake leaves caused by the Xoo strains as indicated below each leaf. Arrow indicates the edge of lesion.

### Selective cloning and function test of TALe genes from the African strain AXO1947

AXO1947 has been sequenced previously and has nine TALe genes and no iTALe gene (Huguet-Tapia et al., 2016). To further demonstrate the feasibility of pZW-Gib to selectively clone TALe genes, a sub-library of TALe genes was constructed by ligating the *Sph*I fragments of genomic DNA from AXO1947 into the *Bsm*BI-digested pZW-Gib. All nine TALe genes were retrieved that matched the annotated TALes in AXO1947 (Supplementary Fig. S4B). The nine TALe genes were individually subcloned into pHM1 and mobilized into ME2 for virulence tests. Only the gene corresponding to a homolog of the previously identified major TALe *TalC*, which targets the S gene *SWEET14*, conferred virulence to ME2 in Kitaake and Zhenshan 97 (Supplementary Fig. 4D).

### Improved vector for cloning TALe genes

Certain TALes, e.g., iTal3a and iTal3b in PXO99^A^, require their unique N-terminal and C-terminal domains for functionality. Two deletions within the N-termini of iTal3a and iTal3b coding sequences are required to suppress the *Xa1*-mediated resistance triggered by TALes (Ji et al., 2016). Other TALe genes simply lack either one *Sph*I site in their DNA sequences due to DNA polymorphisms. The vector pZW-Gib is not able to capture such sequences from the genomes. Furthermore, pZW-Gib derived TALe clones need to be moved into a broad range vector, pHM1 in this case, for further characterization of function in Xoo. To overcome such constraints, a cloning vector capable of capturing of the *Bam*HI fragments and avoiding lack of *Sph*I restriction site of natural TALe genes and direct transfer to Xoo was devised, using pHM1 as the recipient vector for TALe *Bam*HI fragments. Differing from pZW-Gib, pHM1-Gib contains the promoter sequence, the 3’ region downstream of the second *Bam*HI site of *pthXo1* from PXO99^A^, and the *ccdB* cassette. Additionally, a sequence of ColE1 (origin of replication from pBluescript) was integrated into pHM1-Gib to increase the copy number of plasmids in *E. coli*, which facilitates DNA isolation and manipulation (Supplementary Fig S1B). For validation, the *Bam*HI fragment of *pthXo2B* was retrieved from the *Bam*HI digestion of PXO61 genomic DNA in pHM1-Gib through Gibson assembly. The resultant plasmids were directly transferred into ME2, and each transformant tested for the ability to cause disease in Kitaake. The newly assembled *pthXo2B* directed induction of *SWEET13*. To further demonstrate the improved capability of pHM1-Gib, all nine TALe genes from two Africa strains CFBP7321 and CFBP7325 were cloned in the vector. Thirty-nine out of 48 clones in CFBP7325 and 45 out of 48 in CFBP7321, respectively, were positive for TALe-containing sequences (Supplementary Fig. S5, S6, S7). A disease assay confirmed the presence of clones for *TalC* and *TalF*, both previously identified major TALe genes, which could direct disease in rice Kitaake (Supplementary Fig. S8).

## DISCUSSION

In present study, we present two complementary vector systems for cloning TALe gene fragments selectively from a genomic pool of fragments through *in vitro* homologous recombination and single strand annealing, a protocol known as the Gibson assembly method. The strategy depends on exonuclease (e.g., T5 exonuclease) to generate the 3’ end overhangs by chewing back the 5’ nucleotides of vector fragments and genomic fragments, and the assembly of vector fragments and selectively the TALe gene fragments in a single reaction. Each vector system has been demonstrated as functional by reproducibly applying the vector to multiple Xoo strains. Approximately 87% (n=122) of the clones are positive for TALe gene fragments, and a total of 44 TALe genes out of 45, with *Tal5b* of PXO61 missing, were retrieved from four Xoo genomes. The results indicate both plasmids are highly efficient and timesaving compared to the conventional cloning methods for cloning of TALe genes from a given *Xanthomonas* genome. The strategy and even the specific vector system are applicable to other *Xanthomonas* species and pathovars (e.g., Xoc) due to the conserved nucleotide sequences around the *Sph*I and *Bam*HI sites of TALe genes. Some species with more divergent TALe sequence, including *X. translucens*, may require customization of the vector annealing regions (Peng et al., 2016).

The prior studies reported the difficult and time-consuming task in cloning a complement of TALe gene fragments from a given genome largely due to a large number of TALe genes that highly conserved in a given genome and the large number of central tandem repeats of individual TALe genes. The cloning was performed by making genomic libraries followed by Southern hybridization- or PCR-based screening, which is laborious and inefficient. Due to such difficulty, only a few from a large number of sequenced and annotated *Xanthomonas* genomes have been subjected to systematic TALe gene cloning and characterization in contrast to (Yang and White, 2004; Cernadas et al., 2014; Yu et al., 2015). For example, Yu et al. pulled out 115 positive clones (representing 18 unique TALe genes) from 3,000 individual transformant colonies (ca. 4%) through screening a library of *Bam*HI fragments of Xoo K74 genome using dot and Southern blotting (Yu et al., 2015). A similar but improved approach was developed to enrich the *Bam*HI fragments of TALe genes from a given genome by applying two additional restriction enzymes (*ApaL*I and *Sfo*I) to further cut the non-TALe gene fragments of genomic DNA. Ligation and screening by PCR yielded 25.0% and 26.9% positive clones of TALe gene fragments for MAI1 and BAI3 strains of Xoo, respectively (Tran et al., 2018a). Both methods require further subcloning of the TALe *Bam*HI fragments from the pUC based vector into TALe gene scaffold and *Xanthomonas*-suitable vector for function analysis.

TALes represent a largest type III effector family of pathogenic bacteria, and some TALes have been found to be essential for bacteria to inflict crop plants (Boch et al., 2010). Such important TALes represent attractive targets to create disease resistance in host crop plants by interfering with the disease mechanism (Schornack et al., 2013). Naturally occurring or gene editing derived alleles with EBE variations, if disruptive to TALe binding, in the promoters of S genes targeted by TALes confer genetically recessive resistance (Chu et al., 2006; Liu et al., 2011; Li et al., 2012; Blanvillain-Baufumé et al., 2017). Multiplex genome editing produced single engineered rice lines that carried multiple mutations in three *SWEET* gene promoters and resulted in broad-spectrum blight resistance (Oliva et al. submitted). Executor R genes along with promoters containing the synthetic EBEs have been used to trap the widely spread cognate TALes for induced resistance (Römer et al., 2009; Hummel et al., 2012). Deployment of such resistances (dominant, recessive, naturally occurring, or artificially made) against TALes create tremendous selection pressure on the dynamic populations of *Xanthomonas* pathogens. The repetitive and multigenic nature provides a repertoire of TALe genes in a given genome to evolve and potentially overcome the resistances. Broad and durable resistance depends on effectively monitoring the evolving pathogenic populations and quickly identifying newly evolving TALes, monitoring tools including a diagnostic kit comprising 7 components based on TALe-mediated disease mechanism (Eom et al. submitted) and the cloning methods as described here. Additionally, the Gibson-based vectors developed here have potential for high throughput profiling of complement TALe genes from multiple *Xanthomonas* strains when bar coded and combined with next generation sequencing technologies, including PacBio and Nanopore platforms. Development of simple, cheap, and efficient cloning system will also advance our understanding of TALe biology.

## EXPERIMENTAL PROCEDURES

### Plant materials, bacterial strains and growth conditions

Rice (*Oryza sativa*) cultivar Kitaake and Zhenshan 97 were used in this study. Plasmids, Xoo, and *Escherichia coli* strains used in this study are listed in Supplementary Table S2. Rice plants were grown in growth chamber with the temperature of 28 °C, relative humidity of 85% and a photoperiod of 12 hours. *E. coli* DB3.0 and Trans1-T1 were grown in Luria-Bertani medium at 37 °C. Xoo strains were grown in either NB (nutrient broth: beef extract, 3 g/L; peptone, 5 g/L) or TSA (tryptone sucrose medium: tryptone, 10 g/L; sucrose, 10 g/L; glutamic acid, 1 g/L) at 28 °C. Antibiotics used in this study include spectinomycin (100 mg/L) and ampicillin (100 mg/L) for bacterial strains containing appropriate plasmids.

### DNA manipulation and plasmid construction

DNA manipulation and PCR were conducted according to standard protocols (Ausubel et al., 1998). Genomic DNA from Xoo was extracted from bacterial cell grown in NB medium using the MagAttract HMW DNA kit (Qiagen, Hilden, Germany) following the manufacturer’s manual. Plasmids were introduced by electroporation into *X. oryzae* and *E. coli* bacterial competent cells as described previously (Yang and White, 2004). Primers for PCR were synthesized by Integrated DNA Technologies (Coralville, IA, USA); PCR was performed using Taq DNA polymerase or Phusion High-Fidelity DNA polymerase (New England BioLabs, Ipswich, MA, USA).

#### Construction of pZW-Gib

The central repeat region of *pthXo1*was replaced by a gBlock, which contain *ccdB* sequence, as following. First the 3’ fragment of a TAL effector gene was PCR-amplified by using the primers TalAatII-F and TalFlH3-R along the genomic DNA of PXO99^A^. The PCR amplicon was cloned into pZW-*pthXo1* at *Aat*II and *Hin*dIII sites through Gibson assembly. The ligation reaction was transferred into *E. coli* Trans1-T1 cells. The positive bacterial colonies were screened by colony PCR basis on primer TalAatII-F and TALFLH3-R. Candidate clones were sequenced for confirmation of *PthXo1* downstream sequence. A gBlock fragment (Supplementary. Fig. S1A) synthesized by Integrated DNA Technologies (Coralville, IA) was cloned into the new pZW-*pthXo1* at *Sph*I sites by Gibson cloning. The ligation reaction was transferred into *E. coil* DB3.0 cells. Candidate clones were sequenced for confirmation of gblock sequence.

#### Construct pZW-Gib-avrXa7 via Gibson cloning

The central repeat region of *avrXa7* was obtained by digesting pZW-avrXa7 with *Sph*I and selecting DNA fragment of right size through electrophoresis in 1% agarose gel. The *Sph*I fragment was cloned into the backbone of pZW-Gib that was derived from *Bsm*BI-digestion and purification from 1% agarose gel. The ligation was transferred into *E. coli* Trans T-1 cells; positive colonies was first screened via PCR using primer P-F and P-R, *Sph*I digestion and eventually sequencing for confirmation of *avrXa7* insertion. Primer sequences are provided in Supplementary Table S3.

#### Construct TALe gene sublibraries of PXO61 and AXO1947

genomic DNA of PXO61 and AXO1947 was digested with SphI and the fragments were separated through electrophoresis in 1% agarose gel. DNA fragments of about 2-5 kb were extracted and purified from the agarose gel and subcloned through Gibson assembly into the backbone of pZW-Gib that was *Bsm*BI-digested and gel-purified. The Gibson Assembly Cloning kit (New England Biolabs, Ipswich, MA) was used by following the manufacturer’s manual. The subsequent process was identical to the construction of pZW-Gib-avrXa7. Finally, a select set of clones were chosen based on the expected sizes of repeats as determined by *Sph*I digestion and were sequenced using primers Tal-*Sph*I-F and P-R.

#### Transform X. oryzae cells with plasmid DNA

Competent cells of Xoo ME2 were prepared and transformation was performed as previously described (Ji et al., 2016). Briefly, TSA+NB medium (nutrient broth, 8 g/L; tryptone, 10 g/L; sucrose, 10 g/L; glutamic acid, 1 g/L) was used to grow *Xoo* cells. An aliquot of 50 µl of bacterial competent cells was mixed with 10 ng (0.5 µl) of plasmid DNA for electroporation using a Bio-Rad electroporation instrument (Bio-Rad, Hercules, CA, USA). An electric field of 15 kV/cm with a resistance of 200 Ω and a capacitance of 25 µF was applied. After pulse delivery, cells were immediately transferred into 1 ml of SOC medium (20 g tryptone, 5 g yeast extract, 4.8 g MgSO4, 3.6 g dextrose, 0.5 g NaCl and 0.2 g KCl per liter) in a 2 ml round-bottom polypropylene tube. After incubation at 28 °C with constant shaking for 2-4 hours, the electroporated cells were plated onto TSA-NB medium containing the appropriate antibiotics and incubated at 28 °C for 3-5 days. Non-electroporated competent cells were used as a control. Positive clones were picked for further analysis.

### Disease assays

Disease assays to test the function of cloned TALe genes were conducted as following. Briefly, Xoo strains were grown in TSA+NB with appropriate antibiotics at 28 °C. Bacterial cells were collected from culture through low-speed (2,300 g) centrifugation, washed twice and suspended in sterile water. The suspensions were adjusted to an optical density of 0.5 at 600 nm (OD600) and were used to inoculate into fully expanded leaves of 7-8 weeks old rice plants using the leaf-tip clipping method (Kauffman et al., 1973). Lesion lengths were measured about 12 to 14 days post inoculation. The disease assays were performed independently at least twice and on at least four plants each time. One-way analysis of variance statistical analyses was performed on all measurements. The Tukey’s honestly significant difference test was used for post analysis of variance pairwise tests for significance, set at 5% (*P*<0.05).

### Sequence analysis of TALe genes in sequenced *Xanthomonas* genomes

Genomes of *Xanthomonas* strains were obtained from the NCBI under accession numbers: Xoo OS198: CP031461.1, Xoo JL25: CP031457.1, Xoo PX086: CP031463.1, Xoo PX079: CP031462.1, Xoo YC11: CP031464.1, Xoo JL33: CP031459.1, Xoo JP01: CP031460.1, Xoo HuN37: CP031456.1, Xoo JL28: CP031458.1, Xoo BAI3: CP025610.1, Xoo MAI1: CP025609.1, Xoo MAI145: CP019092.1, Xoo MAI134: CP019091.1, Xoo MAI129: CP019090.1, Xoo MAI106: CP019089.1, Xoo MAI99: CP019088.1, Xoo MAI95: CP019087.1, Xoo MAI73: CP019086.1, Xoo MAI68: CP019085.1, Xoo PXO61: CP021789.1, Xoo PXO145: CP013961.1, Xoo AXO1947: CP013666.1, Xoo XF89b: CP011532.1, Xoo PXO602: CP013679.1, Xoo PXO563: CP013678.1, Xoo PXO524: CP013677.1, Xoo PXO282: CP013676.1, Xoo PXO236: CP013675.1, Xoo PXO211: CP013674.1, Xoo PXO71: CP013670.1, Xoo PXO83: CP012947.1, Xoo PXO99A: CP000967.2, Xoo PXO86: CP007166.1, Xoo MAFF 311018: NC_007705.1, Xoo KACC 10331: AE013598.1, Xoc CFBP7331: CP011958.1, Xoc CFBP2286: CP011962.1, Xoc RS105: CP011961.1, Xoc L8: CP011960.1, Xoc CFBP7341: CP011959.1, Xoc BXOR1: CP011957.1, Xoc BLS279: CP011956.1, Xoc B8-12: CP011955.1, Xoc CFBP7342: CP007221.1, Xoc BLS256: CP003057.2, Xcv CFBP7113: CP022270.1, Xcv CFBP7112: CP022269.1, Xcm XcmH1005: CP013004.1.

Analysis of all TALe gene sequences was performed through NCBI blast and further confirmed by using SnapGene software. Sequences used for comparison are 5’-GCATGCATGGCGCAATGCACTGACGGGTGC-3’ and 5’-ACGCCGGATCAGGCGTCTTTGCATGC-3’ around the two *Sph*I restriction sites; while another pair of sequences 5’-GGATCCCATTCGTTCGCGCACGCCAAGTCCTGCCCGCG-3’ and 5’-ACCAGGATCGGGGGCGGCCTCCCGGATCC-3’ around the two *Bam*HI sites of TALe genes.

## ACKNOWLEDGEMENTS

The work was partially supported by subawards to MU and UF from Heinrich Heine University of Dusseldorf, which was funded by the Bill & Melinda Gates Foundation [OPP1155704] (B.Y., F.F.W), the China Scholar Council (C.L., as a point PhD student). The authors are grateful to Wolf F. Frommer for critical read and constructive comments to the manuscript.

