## Supplementary Tables and Figures for "An efficient method to clone TAL effector genes from *Xanthomonas oryzae* using Gibson assembly"

Table S1 to S3

Fig. S1 to S8

**Supplementary Table 1.** Number of TALe genes from different *Xanthomonas* that can be selectively cloned with Gibson assembly.

| *Xanthomonas* | | | Number of TALe genes in genome | Number of TALe genes clonable as *Bam*HI-digested fragments | Number of TALe genes clonable as *Sph*I-digested fragments |
|---|---|---|---|---|---|
| **Strain** | | **Accession No.** | | | |
| ***X. o.* pv. oryzae** (Asia) | **PXO61** | CP033187 | 18 | 17 | 17 |
| | **XF89b** | GCA_002023005.1 | 17 | 17 | 17 |
| | **PXO602** | GCA_001746735.1 | 20 | 19 | 20 |
| | **PXO563** | GCA_001746715.1 | 18 | 17 | 17 |
| | **PXO524** | GCA_001746695.1 | 19 | 17 | 18 |
| | **PXO282** | GCA_001746675.1 | 18 | 17 | 17 |
| | **PXO236** | GCA_001746655.1 | 16 | 14 | 15 |
| | **PXO211** | GCA_001746635.1 | 17 | 15 | 16 |
| | **PXO145** | GCA_001746615.1 | 18 | 15 | 17 |
| | **PXO71** | GCA_001746595.1 | 20 | 17 | 18 |
| | **PXO83** | GCA_001518895.1 | 18 | 15 | 17 |
| | **PXO99**[A] | GCA_000019585.2 | 19 | 18 | 18 |
| | **PXO86** | GCA_000948075.1 | 18 | 15 | 17 |
| | **MAFF311018** | GCA_000010025.1 | 17 | 16 | 16 |
| | **KACC10331** | AE013598.1 | 13 | 13 | 13 |
| | **HuN37** | GCA_003382775.1 | 18 | 17 | 17 |
| | **JL25** | GCA_003382795.1 | 16 | 15 | 15 |
| | **JL28** | GCA_003382815.1 | 12 | 10 | 10 |
| | **JL33** | GCA_003382835.1 | 16 | 15 | 15 |
| | **JP01** | GCA_003382855.1 | 17 | 16 | 16 |
| | **OS198** | GCA_003382875.1 | 18 | 18 | 18 |
| | **YC11** | GCA_003382935.1 | 12 | 12 | 12 |
| ***X. o.* pv. oryzae** (Africa) | **BAI3 (CFBP7321)** | GCA_003031385.1 | 9 | 9 | 9 |
| | **MAI1 (CFBP7325)** | GCA_003031365.1 | 9 | 9 | 8 |
| | **MAI145** | GCA_002850095.1 | 9 | 9 | 8 |
| | **MAI134** | GCA_002850175.1 | 9 | 9 | 7 |
| | **MAI129** | GCA_002850155.1 | 9 | 9 | 8 |
| | **MAI106** | GCA_002850135.1 | 9 | 9 | 8 |
| | **MAI99** | GCA_002850215.1 | 9 | 9 | 8 |
| | **MAI95** | GCA_002850195.1 | 9 | 9 | 8 |
| | **MAI73** | GCA_002850075.1 | 9 | 9 | 8 |
| | **MAI68** | GCA_002850115.1 | 9 | 9 | 8 |
| | **AXO1947** | GCA_001466505.1 | 9 | 9 | 9 |
| ***X. o.* pv. oryzicola** | **CFBP2286** | GCA_001042735.1 | 28 | 27 | 22 |
| | **B8-12** | GCA_001042745.1 | 28 | 27 | 25 |
| | **BLS256** | GCA_000168315.3 | 28 | 27 | 23 |
| | **BLS279** | GCA_001042775.1 | 26 | 25 | 23 |
| | **BXOR1** | GCA_001042795.1 | 25 | 24 | 25 |
| | **CFBP7331** | GCA_001042815.1 | 20 | 20 | 19 |
| | **CFBP7341** | GCA_001042835.1 | 20 | 19 | 18 |
| | **CFBP7342** | GCA_000940825.1 | 23 | 23 | 21 |
| | **L8** | GCA_001042855.1 | 29 | 28 | 26 |
| | **RS105** | GCA_001042875.1 | 24 | 23 | 22 |

| | | | | | |
|---|---|---|---|---|---|
| ***X. citri* pv. vignicola** | **CFBP7112** | GCA_002218265.1 | 1 | 0 | 1 |
| | **CFBP7113** | GCA_002218285.1 | 1 | 0 | 1 |
| ***X. citri* pv. malvacearum** | **XCMH1005** | GCA_002224525.1 | 6 | 6 | 5 |
| **Total TALe genes** | | | 733 | 693 (94%) | 676 (92%) |

**Supplementary Table 2**. Plasmids and bacterial strains used in this study

| **Designation** | **Genotypes or related characteristics** | **Source or reference** |
|---|---|---|
| **Plasmids** | | |
| pHM1 | Broad host range, resistance to spectinomycin, *cos* site | (Hopkins *et al.*, 1992) |
| pZWpthXo1 | *Bam*HI fragment of *pthXo1* in pZW | (Yang & White, 2004) |
| pZW-Gib | *Sph*I fragment of gblock in pZWpthXo1 | This study |
| pZWpxo61tale1a | *SphI* fragment of *PXO61tale1a* in pZW-Gib | This study |
| pZWpxo61tale1b | *Sph*I fragment of *PXO61tale1b* in pZW-Gib | This study |
| pZWpxo61tale1c | *Sph*I fragment of *PXO61tale1c* in pZW-Gib | This study |
| pZWpxo61tale1d | *Sph*I fragment of *PXO61tale1d* in pZW-Gib | This study |
| pZWpxo61tale2a | SphI fragment of *PXO61tale2a* in pZW-Gib | This study |
| pZWpxo61tale2b | *Sph*I fragment of *PXO61tale2b* in pZW-Gib | This study |
| pZWpxo61tale2c | *Sph*I fragment of *PXO61tale2c* in pZW-Gib | This study |
| pZWpxo61tale3a | *Sph*I fragment of *PXO61tale3a* in pZW-Gib | This study |
| pZWpxo61tale3b | *Sph*I fragment of *PXO61tale3b* in pZW-Gib | This study |
| pZWpxo61tale4a | *Sph*I fragment of *PXO61tale4a* in pZW-Gib | This study |
| pZWpxo61tale4b | *Sph*I fragment of *PXO61tale4b* in pZW-Gib | This study |
| pZWpxo61tale4c | *Sph*I fragment of *PXO61tale4c* in pZW-Gib | This study |
| pZWpxo61tale5a | *Sph*I fragment of *PXO61tale5a* in pZW-Gib | This study |
| pZWpxo61tale6a | *Sph*I fragment of *PXO61tale6a* in pZW-Gib | This study |
| pZWpxo61tale6b | *Sph*I fragment of *PXO61tale6b* in pZW-Gib | This study |
| pZWpxo61tale6c | *Sph*I fragment of *PXO61tale6c* in pZW-Gib | This study |
| pZWpxo61tale7 | *Sph*I fragment of *PXO61tale7* in pZW-Gib | This study |
| pZWaxo1947tale1 | *Sph*I fragment of *AXO1947tale1* in pZW-Gib | This study |
| pZWaxo1947tale2 | *Sph*I fragment of *AXO1947tale2* in pZW-Gib | This study |
| pZWaxo1947tale3 | *Sph*I fragment of *AXO1947tale3* in pZW-Gib | This study |
| pZWaxo1947tale4a | *Sph*I fragment of *AXO1947tale4a* in pZW-Gib | This study |
| pZWaxo1947tale4b | *Sph*I fragment of *AXO1947tale4b* in pZW-Gib | This study |
| pZWaxo1947tale4c | *Sph*I fragment of *AXO1947tale4c* in pZW-Gib | This study |
| pZWaxo1947tale5 | *Sph*I fragment of *AXO1947tale5* in pZW-Gib | This study |
| pZWaxo1947tale6 | *Sph*I fragment of *AXO1947tale6* in pZW-Gib | This study |
| pZWaxo1947tale7 | *Sph*I fragment of *AXO1947tale7* in pZW-Gib | This study |
| pHM1-Gib | pHM1 vector with natural TALe's promoter and colE1replication origin, *Bam*HI fragment of gblock | This study |

| | | |
|---|---|---|
| pHM1-pZWpxo61tale1a | pZWpxo61tale1a in pHM1 | This study |
| pHM1-pZWpxo61tale1b | pZWpxo61tale1b in pHM1 | This study |
| pHM1-pZWpxo61tale1c | pZWpxo61tale1c in pHM1 | This study |
| pHM1-pZWpxo61tale1d | pZWpxo61tale1d in pHM1 | This study |
| pHM1-pZWpxo61tale2a | pZWpxo61tale2a in pHM1 | This study |
| pHM1-pZWpxo61tale2b | pZWpxo61tale2b in pHM1 | This study |
| pHM1-pZWpxo61tale2c | pZWpxo61tale2c in pHM1 | This study |
| pHM1-pZWpxo61tale3a | pZWpxo61tale3a in pHM1 | This study |
| pHM1-pZWpxo61tale3b | pZWpxo61tale3b in pHM1 | This study |
| pHM1-pZWpxo61tale4a | pZWpxo61tale4a in pHM1 | This study |
| pHM1-pZWpxo61tale4b | pZWpxo61tale4b in pHM1 | This study |
| pHM1-pZWpxo61tale4c | pZWpxo61tale4c in pHM1 | This study |
| pHM1-pZWpxo61tale5a | pZWpxo61tale5a in pHM1 | This study |
| pHM1-pZWpxo61tale6a | pZWpxo61tale6a in pHM1 | This study |
| pHM1-pZWpxo61tale6b | pZWpxo61tale6b in pHM1 | This study |
| pHM1-pZWpxo61tale6c | pZWpxo61tale6c in pHM1 | This study |
| pHM1-pZWpxo61tale7 | pZWpxo61tale7 in pHM1 | This study |
| pHM1-pZWaxo1947tale1 | pZWaxo1947tale1 in pHM1 | This study |
| pHM1-pZWaxo1947tale2 | pZWaxo1947tale2 in pHM1 | This study |
| pHM1-pZWaxo1947tale3 | pZWaxo1947tale3 in pHM1 | This study |
| pHM1-pZWaxo1947tale4a | pZWaxo1947tale4a in pHM1 | This study |
| pHM1-pZWaxo1947tale4b | pZWaxo1947tale4b in pHM1 | This study |
| pHM1-pZWaxo1947tale4c | pZWaxo1947tale4c in pHM1 | This study |
| pHM1-pZWaxo1947tale5 | pZWaxo1947tale5 in pHM1 | This study |
| pHM1-pZWaxo1947tale6 | pZWaxo1947tale6 in pHM1 | This study |
| pHM1-pZWaxo1947tale7 | pZWaxo1947tale7 in pHM1 | This study |
| pHM1-Gib-CFBP7321-TalC | BamHI fragment of *CFBP7321talc* in pHM1-Gib | This study |
| pHM1-Gib-CFBP7325-TalF | BamHI fragment of *CFBP7325talf* in pHM1-Gib | This study |

**Bacterial Strains**

*Escherichia coli*

| | | |
|---|---|---|
| T1 | F-φ80(lacZ)ΔM15ΔlacX74hsdR(rk$^-$, mk$^+$)ΔrecA1398endA1tonA | Thermo Fisher Scientific |
| DB3.0 | Str$^R$, *gyrA*462 *endA*1 Δ*(sr*1-*rec*A*)* *mcrB mrr hsdS*20 *glnV*44 *ara*14 *galK*2 *lac*Y1 | (Bernard & Couturier, 1992) |

| | |
|---|---|
| | *proA*2 *rps*L20 *xyl*5 *leuB*6 *mtl*1, resistant to ccdB |

*Xanthomonas oryzae* pv. *oryzae*

| | | |
|---|---|---|
| PXO99$^A$ | Philippine race 6, azacytidine resistant clone of PXO99 | (Hopkins et al., 1992) |
| PXO61 | Philippine race 2 | (Yang & White, 2004) |
| AXO1947 | African strain | (Huguet-Tapia *et al.*, 2016) |
| ME2 | Mutant of PXO99$^A$ with *pthXo1* inactivated | (Yang & White, 2004) |
| ME2(pHM1) | PXO99$^A$ with pBY1 inserted into pthXo1 | (Yang & White, 2004) |
| ME2 (pHM1pZWpxo61tale1a) | ME2 with pHM1-pZWpxo61tale1a | This study |
| ME2(pHM1-pZWpxo61tale1b) | ME2 with pHM1-pZWpxo61tale1b | This study |
| ME2(pHM1-pZWpxo61tale1c) | ME2 with pHM1-pZWpxo61tale1c | This study |
| ME2(pHM1-pZWpxo61tale1d) | ME2 with pHM1-pZWpxo61tale1d | This study |
| ME2(pHM1-pZWpxo61tale2a) | ME2 with pHM1-pZWpxo61tale2a | This study |
| ME2(pHM1-pZWpxo61tale2b) | ME2 with pHM1-pZWpxo61tale2b | This study |
| ME2(pHM1-pZWpxo61tale2c) | ME2 with pHM1-pZWpxo61tale2c | This study |
| ME2(pHM1-pZWpxo61tale3a) | ME2 with pHM1-pZWpxo61tale3a | This study |
| ME2(pHM1-pZWpxo61tale3b) | ME2 with pHM1-pZWpxo61tale3b | This study |
| ME2(pHM1-pZWpxo61tale4a) | ME2 with pHM1-pZWpxo61tale4a | This study |
| ME2(pHM1-pZWpxo61tale4b) | ME2 with pHM1-pZWpxo61tale4b | This study |
| ME2(pHM1-pZWpxo61tale4c) | ME2 with pHM1-pZWpxo61tale4c | This study |
| ME2(pHM1-pZWpxo61tale5a) | ME2 with pHM1-pZWpxo61tale5a | This study |
| ME2(pHM1-pZWpxo61tale6a) | ME2 with pHM1-pZWpxo61tale6a | This study |
| ME2(pHM1-pZWpxo61tale6b) | ME2 with pHM1-pZWpxo61tale6b | This study |
| ME2(pHM1-pZWpxo61tale6c) | ME2 with pHM1-pZWpxo61tale6c | This study |
| ME2(pHM1-pZWpxo61tale7) | ME2 with pHM1-pZWpxo61tale7 | This study |
| ME2(pHM1-pZWaxo1947tale1) | ME2 with pHM1-pZWaxo1947tale1 | This study |

| | | |
|---|---|---|
| ME2(pHM1-pZWaxo1947tale2) | ME2 with pHM1-pZWaxo1947tale2 | This study |
| ME2(pHM1-pZWaxo1947tale3) | ME2 with pHM1-pZWaxo1947tale3 | This study |
| ME2(pHM1-pZWaxo1947tale4a) | ME2 with pHM1-pZWaxo1947tale4a | This study |
| ME2(pHM1-pZWaxo1947tale4b) | ME2 with pHM1-pZWaxo1947tale4b | This study |
| ME2(pHM1-pZWaxo1947tale4c) | ME2 with pHM1-pZWaxo1947tale4c | This study |
| ME2(pHM1-pZWaxo1947tale5) | ME2 with pHM1-pZWaxo1947tale5 | This study |
| ME2(pHM1-pZWaxo1947tale6) | ME2 with pHM1-pZWaxo1947tale6 | This study |
| ME2(pHM1-pZWaxo1947tale7) | ME2 with pHM1-pZWaxo1947tale7 | This study |
| ME2(pHM1-CFBP7321-TalC) | ME2 with pHM1-CFBP7321-TalC | This study |
| ME2(pHM1-CFBP7325-TalF) | ME2 with pHM1-CFBP7325-TalF | This study |

**Supplementary Table 3.** Primers used in this study

| Name | Sequence (5' to 3') | Usage |
|---|---|---|
| TalAatII-F | TTGGCCTGCCTCGGCGGACGTCCT | Amplify the 3' region of TALe gene |
| TalFlH3-R | TGGCGGCCGCTCTAGGCCaagcttTCActtatcgtcatcgtccttgtaatcGATCGTCCCTCCGACTGAG | |
| P-F | AGAGCATTGTTGCCCAGT | PCR screening clones for TALe genes |
| P-R | TCTGATCTCCCTCGTGCATTG | |
| Tal-SphI-F | AGTTGGACACAGGCCAACTTC | Sequencing TALe repeat region from the 5' end |

| | | |
|---|---|---|
| P-R | TCTGATCTCCCTCGTGCATTG | Sequencing TALe repeat region from the 3’ end |

**A**

**gBlock1 sequence for pZW-Gib**

*Sph*I *Bsm*BI
GGCGTGACCGCAGTGGAGGCAGT***GCATGC***ATGGCGCAATGCACTGACGGGTGCA***GAGACG***ATAA
CAGTATGCGTATTTGCGCGCTGATTTTTGCGGTATAAGAATATATACTGATATGTATACCCGAA
GTATGTCAAAAAGAGGTATGCTATGAAGCAGCGTATTACAGTGACAGTTGACAGCGACAGCTAT
CAGTTGCTCAAGGCATATATGATGTCAATATCTCCGGTCTGGTAAGCACAACCATGCAGAATGA
AGCCCGTCGTCTGCGTGCCGAACGCTGGAAAGCGGAAAATCAGGAAGGGATGGCTGAGGTCGCC
CGGTTTATTGAAATGAACGGCTCTTTTGCTGACGAGAACAGGGGCTGGTGAAATGCAGTTTAAG
GTTTACACCTATAAAAGAGAGAGCCGTTATCGTCTGTTTGTGGATGTACAGAGTGATATTATTG
ACACGCCCGGGCGACGGATGGTGATCCCCCTGGCCAGTGCACGTCTGCTGTCAGATAAAGTCTC
CCGTGAACTTTACCCGGTGGTGCATATCGGGGATGAAAGCTGGCGCATGATGACCACCGATATG
GCCAGTGTGCCGGTCTCCGTTATCGGGGAAGAAGTGGCTGATCTCAGCCACCGCGAAAATGACA
TCAAAAACGCCATTAACCTGATGTTCTGGGGAATATAATGAGGCTCCCTTATACAC***CGTCTC***AA
*Bsm*BI
CGCCGGATCAGGCGTCTTT***GCATGC***ATTCGCCGATTCGCTGGAG
*Sph*I

**B**

**gBlock2 sequence for pHM1-Gib**:

*Hind*II
CACGCCAAGTCCTGCCCGCG***AAGCTT***GCATTAGGCACCCCAGGCTTTACACTTTATGCTTCCGG
CTCGTATAATGTGTGGATTTTGAGTTAGGATCGATCCGGCTTACTAAAAGCCAGATAACAGTAT
GCGTATTTGCGCGCTGATTTTTGCGGTATAAGAATATATACTGATATGTATACCCGAAGTATGT
CAAAAAGAGGTATGCTATGAAGCAGCGTATTACAGTGACAGTTGACAGCGACAGCTATCAGTTG
CTCAAGGCATATATGATGTCAATATCTCCGGTCTGGTAAGCACAACCATGCAGAATGAAGCCCG
TCGTCTGCGTGCCGAACGCTGGAAAGCGGAAAATCAGGAAGGGATGGCTGAGGTCGCCCGGTTT
ATTGAAATGAACGGCTCTTTTGCTGACGAGAACAGGGGCTGGTGAAATGCAGTTTAAGGTTTAC
ACCTATAAAAGAGAGAGCCGTTATCGTCTGTTTGTGGATGTACAGAGTGATATTATTGACACGC
CCGGGCGACGGATGGTGATCCCCCTGGCCAGTGCACGTCTGCTGTCAGATAAAGTCTCCCGTGA
ACTTTACCCGGTGGTGCATATCGGGGATGAAAGCTGGCGCATGATGACCACCGATATGGCCAGT
GTGCCGGTCTCCGTTATCGGGGAAGAAGTGGCTGATCTCAGCCACCGCGAAAATGACATCAAAA
ACGCCATTAACCTGATGTTCTGGGGAATATAAATGTCAGGCTCCCTTATACACAGCCAGTCTGC
AGGTCGA***AAGCTT***ACCAGGATCGGGGGCGGCCT
*Hind*II

**Supplementary Fig. S1.** DNA sequences of gBlock fragments synthesized to make two Gibson cloning vectors. A. gBlock1 was used to inserted into pZW-pthXo1 at *Sph*I sites through Gibson cloning. The two sequences shaded in yellow color are homologous to the ends of *Sph*I fragments of TALe genes. B. gBlock2 was used to make pHM1-Gib. The two sequences shaded in yellow color are homologous to the ends of *Bam*HI fragments of TALe genes.

| $TALE_{AXO1947}$ | RVDs |
|---|---|
| | 1 2 3 4 5 6 7 8 9 10 11 12 13 14 15 16 17 18 19 20 21 22 23 24 25 26 |
| TALE1 | NN-NN-NN-NI-NN-NN-HD-NI-NG-HD-NI-NN-HD-NG-HD-HD-NG |
| TALE2 | NN-NN-NN-HD-NI-NN-HD-HD-HD-NI-NN-NN-HD-HD-N*-NG-HD-NI |
| TALE3 | NI-HD-NN-NS-NN-NG-HD-NG-HD-NG-NN-NG-HD-NS-HD-NI-NG-HD-HD-NN-HD-NN |
| TALE4a | NN-ND-NN-NI-NK-NN-HD-NN-NG-NG-N*-HD-N*-HD-NI-NN-HD-NG-HD-HD-HD-NG-NN-HD-HD-NG |
| TALE4b | NN-HD-NI-NN-HD-NG-HD-HD-NG-NG-NI-NG-NI-NG |
| TALE4c | NN-NG-NN-HD-HD-NI-N*-NG-HD-NI-NG-NN-HD-NI-NG-NI-NG-NN-NG-HD-NI-NI-NG-HD-NN-NG |
| TALE5 | NN-HD-HD-NN-NN-NG-NG-HD-NG-HD-HD-NG-HD-HD-HD-HD-NG-HD-NG |
| TALE6(TalC) | NS-NG-NS-HD-NI-NG-NN-NG-HD-NI-NN-N*-NI-NN-HD-NG-NI-NN-N*-HD-NN-NG |
| TALE7 | NN-HD-NV-HD-NI-NG-NI-NN-NS-HD-HD-NI-NG-NI-NG-NI |

**Supplementary Fig. S2** Nine TALe genes from AXO1947 were cloned using pZW-Gib vector and Gibson assembly method. The RVDs of individual TALes are shown under numbers (1 to 26) indicating the order of 33-34 amino acid repeats. Asterisk (*) indicates amino acid at the $13^{th}$ position missing.

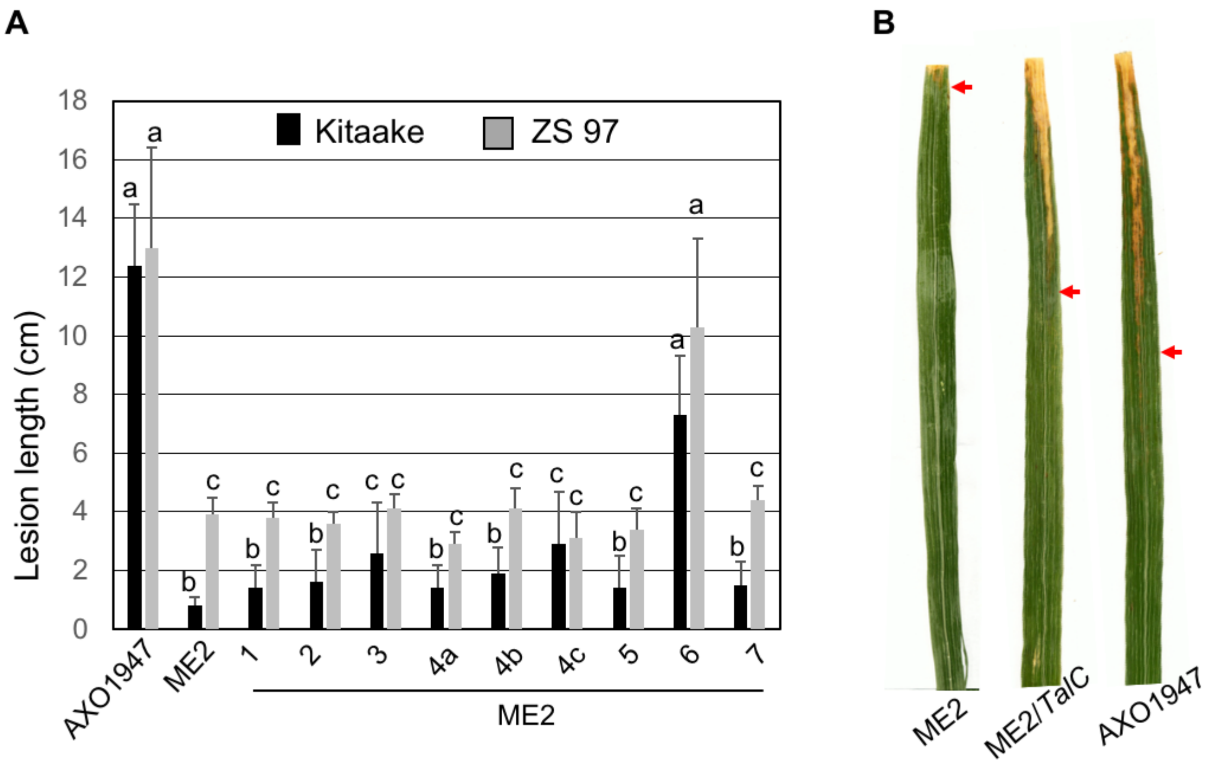

**Supplementary Fig. S3** Virulence contribution of nine TALes cloned from AXO1947. **A.** Lesion lengths caused in rice Kitaake and Zhenshan 97 (ZS 97) by different Xoo strains indicated below the paired columns. Different lower letters indicate statistically significant difference (means ± s.e.m, n=10, $P$<0.05). **B.** blight symptom in Kitaake leaves caused by the Xoo strains as indicated below each leaf. Arrow indicates the edge of lesion.

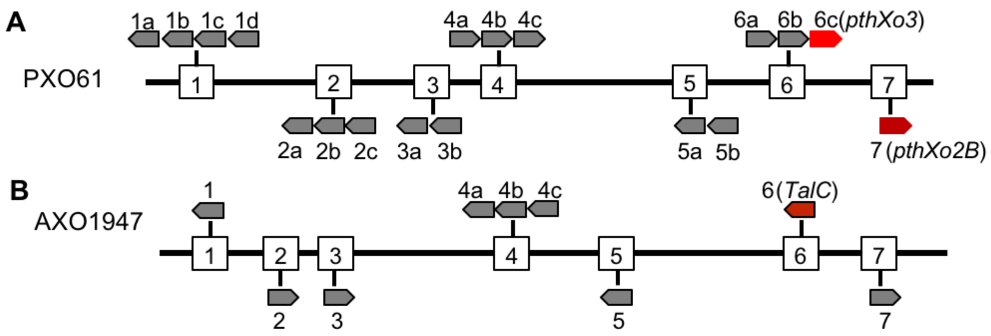

C

| Strain | | Lesion length (cm) |
|---|---|---|
| ME2 | | 2.3±0.3 |
| ME2 | *avrXa7* | 10.2±3.4 |
| | TALE1a-PXO61 | 1.5±0.7 |
| | TALE1b-PXO61 | 1.1±0.7 |
| | TALE1c-PXO61 | 0.5±0.1 |
| | TALE1d-PXO61 | 0.5±0.0 |
| | TALE2a-PXO61 | 0.9±0.4 |
| | TALE2b-PXO61 | 0.6±0.2 |
| | TALE2c-PXO61 | 0.6±0.2 |
| | TALE3a-PXO61 | 1.5±0.7 |
| | TALE3b-PXO61 | 2.2±1.8 |
| | TALE4a-PXO61 | 1.4±0.8 |
| | TALE4b-PXO61 | 1.2±0.8 |
| | TALE4c-PXO61 | 1.0±0.4 |
| | TALE5-PXO61 | 1.3±0.8 |
| | TALE6a-PXO61 | 1.3±0.3 |
| | TALE6b-PXO61 | 1.4±1.1 |
| | TALE6c-PXO61 (*pthXo3*) | 9.6±1.8 |
| | TALE7-PXO61 (*pthXo2B*) | 10.3±3.4 |

D

| Strain | | Lesion length (cm) |
|---|---|---|
| ME2 | TALE1_1947 | 1.4±0.8 |
| | TALE2_1947 | 1.6±1.0 |
| | TALE3_1947 | 2.6±1.7 |
| | TALE4a_1947 | 1.7±0.8 |
| | TALE4b_1947 | 1.9±0.9 |
| | TALE4c_1947 | 1.4±0.8 |
| | TALE5_1947 | 1.4±1.1 |
| | TALE6_1947 (*TalC*) | 5.1±1.4 |
| | TALE7_1947 | 1.5±0.8 |

**ME2** is a PXO99 mutant with the TALE gene *pthXo1* inactivated

**Supplementary Fig. S4** TALe genes from PXO61 and AXO1947. **A.** TALe genes clustered in the genome of PXO61. **B**. TALe genes clustered in the genome of AXO1947. **C**, **D**. Lesion length measurements (mean ± s.e.m, n=10) in Kitaake caused by different strains.

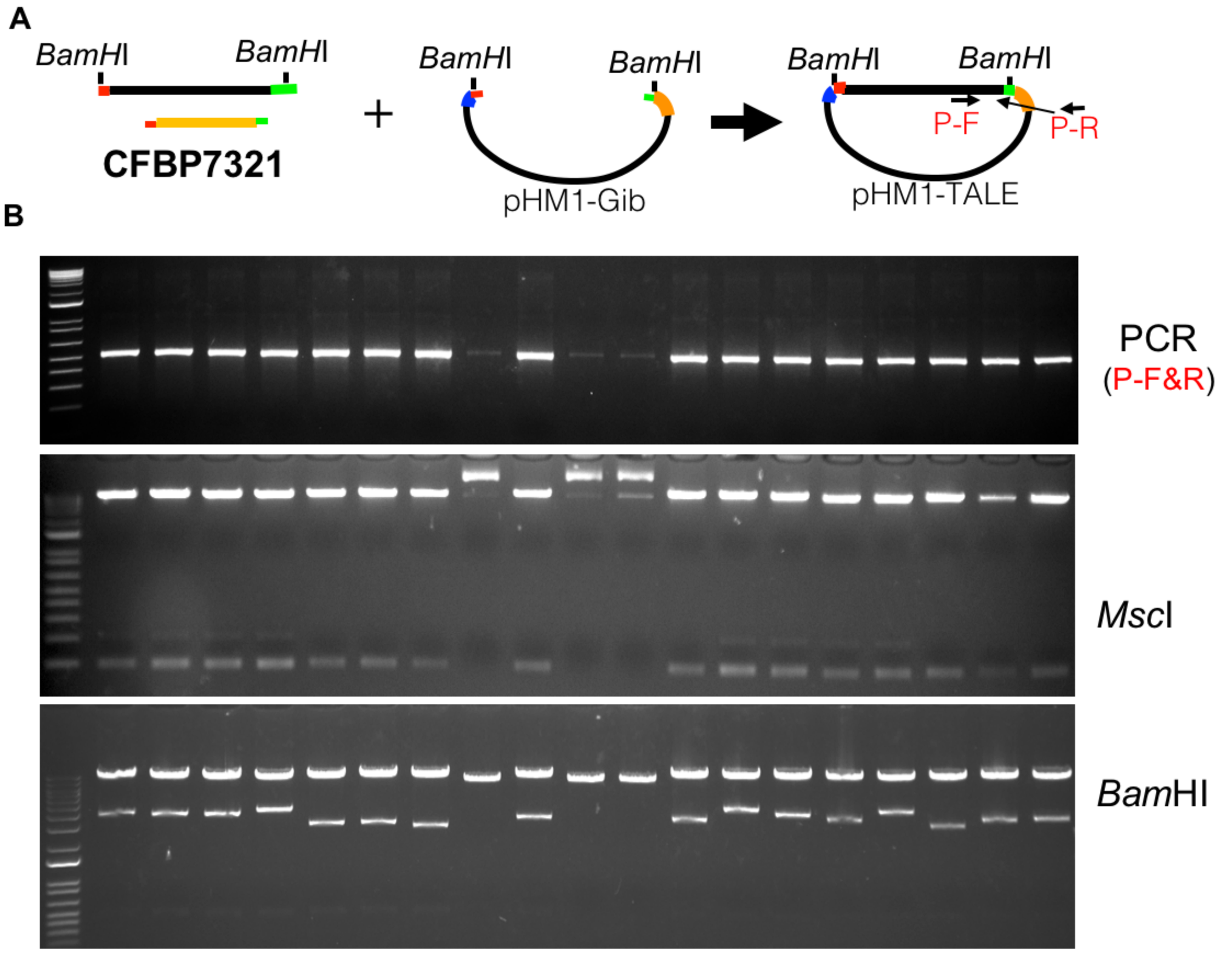

**Supplementary Fig. S5** Validation of TALe clones through PCR, restriction enzyme digestions. **A.** Schematics of selective isolation of *Bam*HI fragments TALe genes from genomic DNA of CFBP7321. **B.** Validation of TALe clones through PCR with primers P-F and P-R of individual clones as indicated above lanes of upper gel image, digestion by *Msc*I which cuts each of central repeats (the DNA band patterns resulted from partial digestion), and digestion by *Bam*HI.

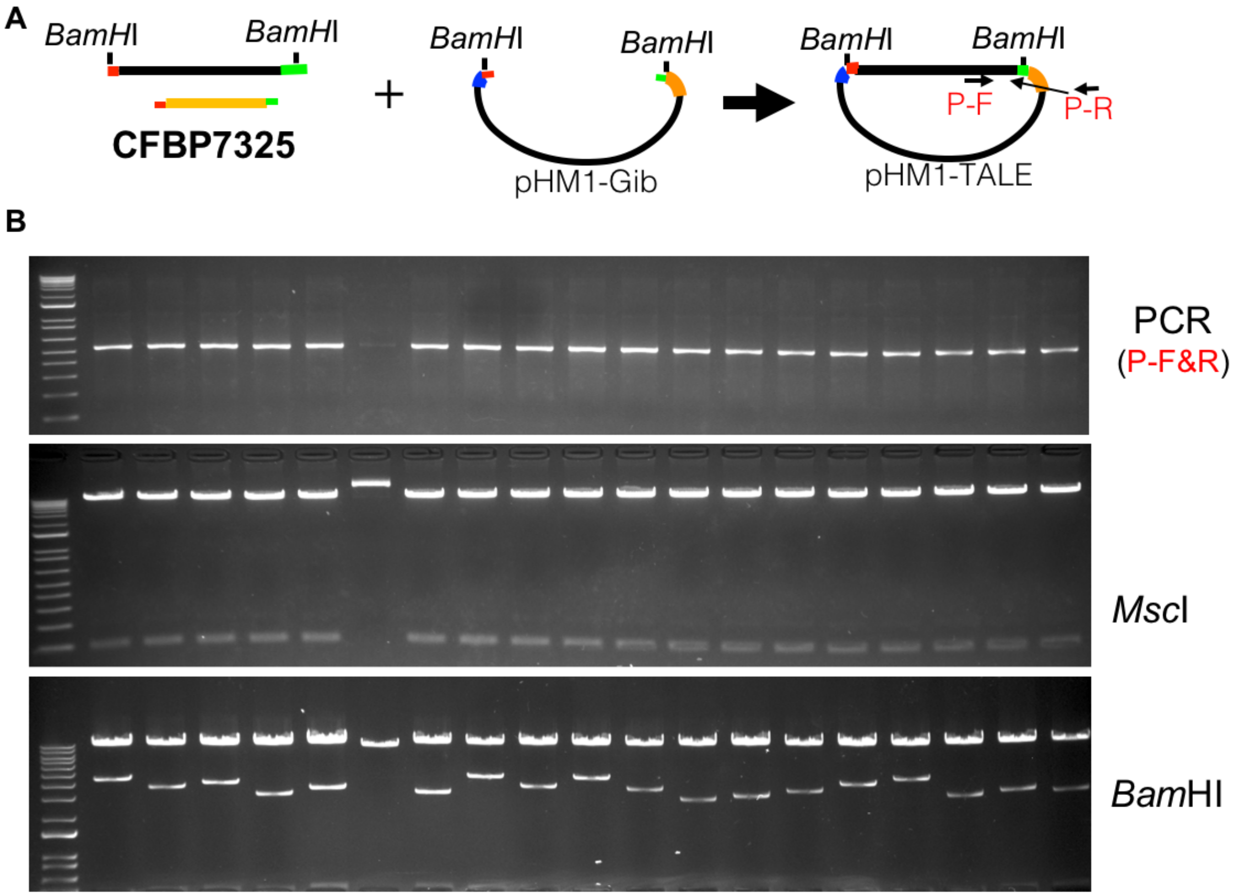

**Supplementary Fig. S6** Validation of TALe clones through PCR, restriction enzyme digestions. **A.** Schematics of selective isolation of *Bam*HI fragments TALe genes from genomic DNA of CFBP7325. **B.** Validation of TALe clones through PCR with primers P-F1 and P-R1 of individual clones as indicated above lanes of upper gel image, digestion by *Msc*I which cuts each of central repeats (the DNA band patterns resulted from partial digestion), and digestion by *Bam*HI.

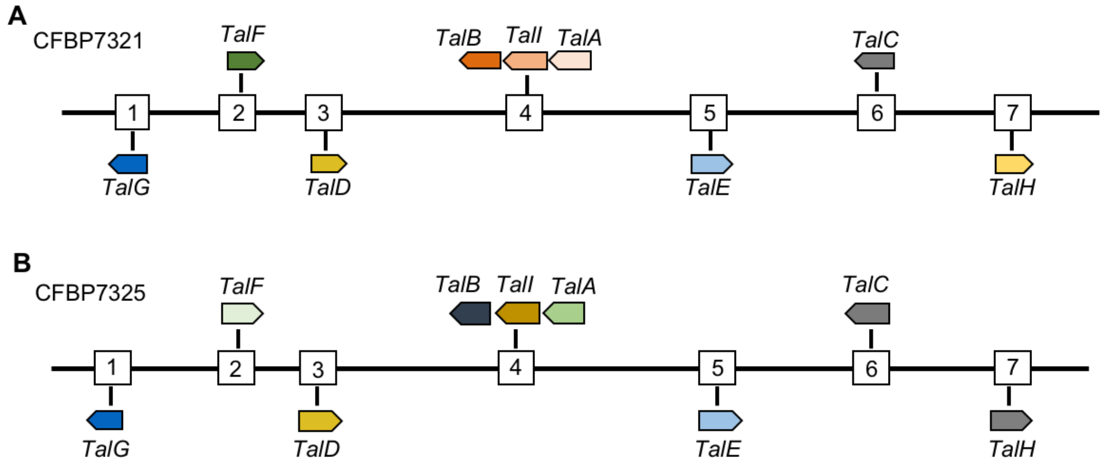

**Supplementary Fig. S7** TALe gene distribution in two African Xoo genomes. **A, B.** Nine TALe genes in each of two Xoo genomes are syntenic in their locations with same colors denoting identical TALes at the amino acid level while different colors indicating different TALes at the amino acid level.

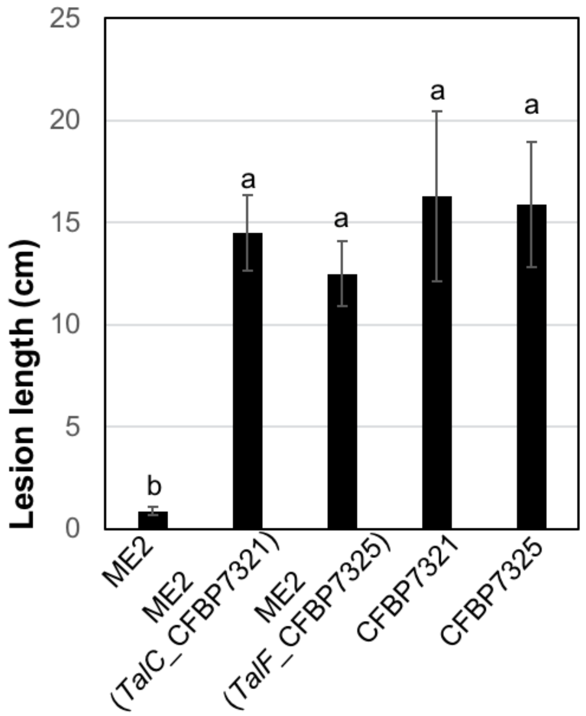

**Supplementary Fig. S8** Two TALe genes cloned with pHM1-Gib system were functional in virulence. The virulence of *TalC* from CFBP7321 and *TalF* from CFBP7325 were tested in Kitaake leaves. Different letters indicate statistically significant difference.
